# Age-dependent VDR peak DNA methylation as a mechanism for latitude-dependent MS risk

**DOI:** 10.1101/2020.04.27.062075

**Authors:** Lawrence T C Ong, Stephen D Schibeci, Nicole L Fewings, David R Booth, Grant P Parnell

## Abstract

**Background:** The mechanisms linking UV radiation and vitamin D exposure to the risk of acquiring the latitude and critical period dependent autoimmune disease, multiple sclerosis, is unclear. We examined the effect of vitamin D on DNA methylation as well as DNA methylation at vitamin D receptor binding sites in adult and paediatric myeloid cells.

**Results:** Very few DNA methylation changes occurred in adult and paediatric cells treated with calcitriol. However, several VDR binding sites across the genome demonstrated increased DNA methylation in cells of adult origin. Genes associated with these VDR binding sites were enriched for intracellular signalling and cell activation pathways, suggesting that age-dependent potential for myeloid cell differentiation and adaptive immune system regulation may be encoded for by DNA methylation.

**Conclusions:** These results suggest vitamin D exposure at critical periods in immune system development may contribute to the well characterised latitude related differences in autoimmune disease incidence.

## Background

The prevalence of autoimmune diseases such as multiple sclerosis (MS), type 1 diabetes mellitus, rheumatoid arthritis and atopic diseases such as asthma follow a latitude gradient, with increasing prevalence at latitudes more distant from the equator^1,2^. Ultraviolet light exposure and hence serum vitamin D levels are known to correlate with latitude, yet a precise mechanism linking vitamin D to immune disease remains elusive. DNA methylation, an important epigenetic mark, has been posited as a potential link between environmental exposures and disease due to its susceptibility to environmental change^3^ and relative stability over time^4^.

Some latitude dependent diseases such as MS also demonstrate a critical period, where risk factors such as latitude of residence appear to exert their influence during childhood and adolescence^5–7^. This critical period is perhaps underpinned by age-related susceptibility to alterations in DNA methylation. DNA methylation changes have been detected in leukocyte development at key histone modifiers, chromatin remodellers and immune susceptibility loci within the first five years of life^8^. DNA methylation changes proceed more rapidly in normal childhood development, with changes in peripheral blood occurring at a three to four-fold higher rate compared with adults^9^. Prenatal susceptibility to environmental insults such as famine, are also highly influenced by gestational age, resulting in persistent DNA methylation changes into adulthood^10,11^.

Vitamin D exerts its genomic effects through binding the vitamin D receptor (VDR). Binding of the active form of vitamin D, calcitriol, results in heterodimerisation of the VDR with the retinoid X receptor (RXR). This heterodimer binds regions of DNA known as vitamin D response elements (VDREs), which lie in the promoter regions of vitamin D responsive genes and lead to subsequent upregulation or suppression of DNA transcription. DNA methylation at VDREs may therefore interfere with the effects of calcitriol on transcriptional regulation.

Whilst the genomic effects of vitamin D have been well characterised through its interactions via the VDR-RXR complex, its effects on DNA methylation have only been partially characterised. A study of human *ex vivo* leukocytes found vitamin D_3_ supplementation during and shortly after pregnancy led to mixed DNA methylation changes in mothers and infants. In infants, methylation loss was associated with genes regulating apoptosis and antigen presentation^12^. In another study, global leukocyte DNA methylation increased in a dose-dependent manner with vitamin D_3_ administration^13^.

There have been few studies on the effect of vitamin D on the methylome of specific immune cell subsets. A study of cholecalciferol supplementation and mouse CD4+ T cells in experimental autoimmune encephalomyelitis (EAE; a mouse model of MS), showed global decreases in DNA methylation. This was associated with changes in the expression of enzymes involved in the establishment and maintenance of DNA methylation marks, with concomitant decreases in CD4+ T cell proliferation and differentiation into inflammatory Th1 and Th17 subsets^14^. Another study found increases in Helios+ Foxp3+ T regulatory cells with 1,25(OH)_2_Vitamin D_3_ (calcitriol) supplementation that were associated with amelioration of EAE, with an increase rather than decrease in global DNA methylation^15^.

More MS risk genes are predominantly expressed in mononuclear phagocytic cells than any other cell subset^16^. These cells are likely to be important in the pathogenesis of MS through their regulation of immune cell differentiation, via mechanisms such as antigen presentation and expression of key vitamin D associated MS risk genes^16,17^. An epigenome wide study of vitamin D treatment on the human monocyte cell line, THP-1, found marked changes in chromatin accessibility due to vitamin D with maximal chromatin opening after 24 hours^18^. Despite this, *ex vivo* mononuclear cells cultured with vitamin D for up to 120 hours did not show any differentially methylated CpGs despite extensive changes in gene expression^19^. The authors suggested donor age may have affected DNA methylation plasticity, however other factors including duration of culture, use of terminally differentiated cells and heterogeneous cell population, may also have contributed to the apparent lack of effect on DNA methylation.

We therefore hypothesised that differentiating haematopoietic progenitors into monocyte/macrophage lineage cells in the presence of sustained calcitriol exposure would result in age-dependent DNA methylation changes. Thus, we sought to determine whether immune cell DNA methylation is affected by exposure to calcitriol. Secondly, because the multiple sclerosis latitude gradient appears to be mediated by a critical period, we sought to determine whether calcitriol-related DNA methylation changes vary with age. Because the genomic effects of vitamin D are mediated by its receptor, we also investigated the DNA methylation changes at corresponding binding sites.

## Results

### Calcitriol Results in Changes in Cell Number, Morphology and Phenotype in Cell Culture

We conducted preliminary cell culture with varying calcitriol concentrations to determine effects on cell morphology and immunophenotype. Using CD34+ haematopoietic progenitor cells originating from an adult subject, at day 22, we noted a marked decrease in cell number and a less activated immunophenotype with higher concentrations of calcitriol (Figure 1). This preliminary experiment confirmed that a physiological concentration of calcitriol (0.1nM) was sufficient to elicit phenotypic changes in cultured cells. In the final experiment that cultured cells from two adult and two paediatric subjects, overall CD14+ percentage as a subset of CD45+ cells, was greater in cells of paediatric origin than those of adult origin with mean CD14+ proportion 78.9% vs 11.7% (p<4.99×10^−4^, two-tailed t-test).

**Figure 1.**
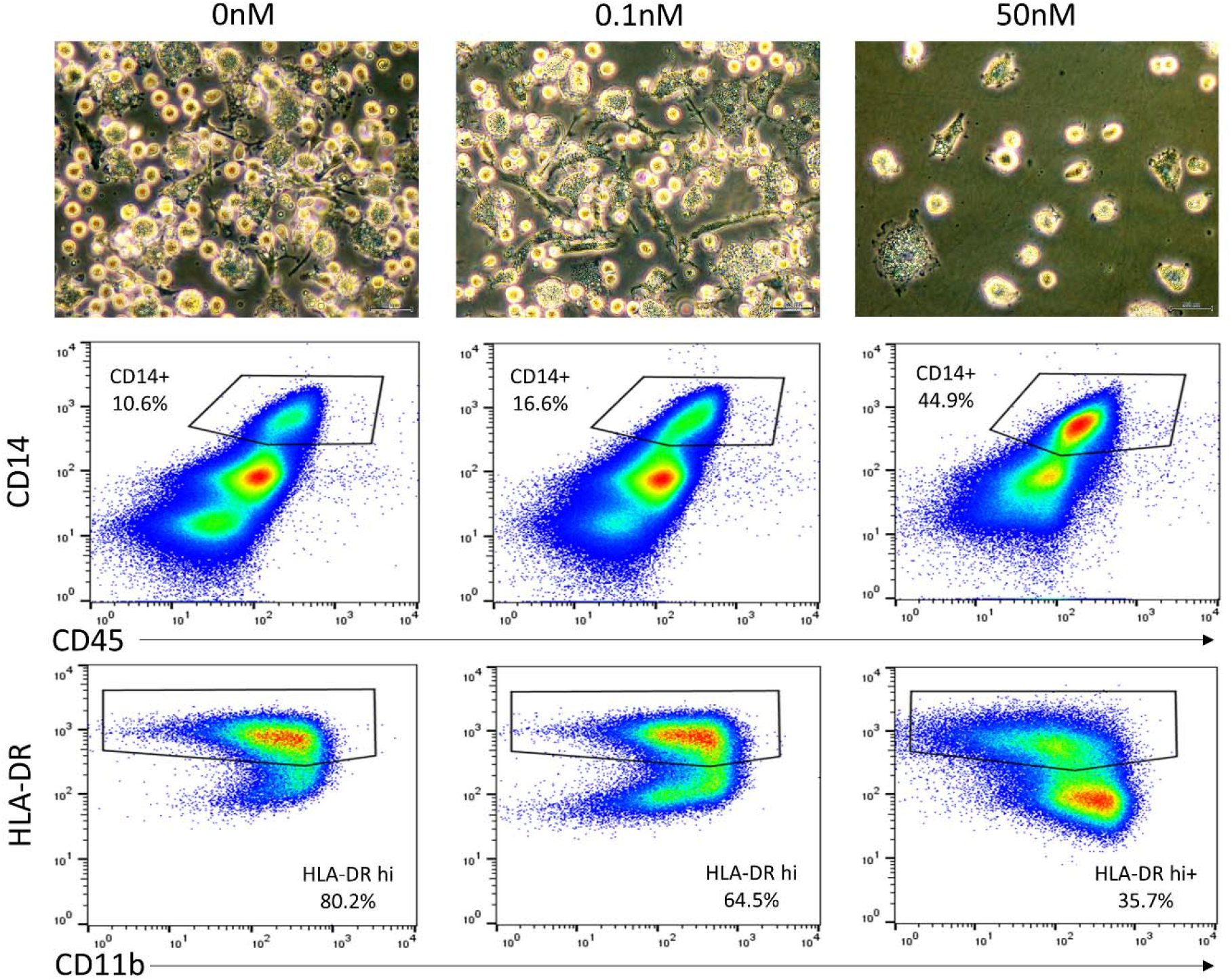
Light microscopy and flow cytometric characterisation of cultured cells at day 22. Cultured cells of adult origin with media containing 0nM calcitriol (left), 0.1nM calcitriol (centre) and 50nM calcitriol (right). There were marked morphologic and immunophenotypic changes, with overall decrease in cell number, fewer fusiform shaped cells, greater CD14+ proportion and decreases in HLA-DR and CD16 expression (not shown) at higher calcitriol concentrations.

### DNA Methylation Varies More Due to Age and Individual Differences Than Due to Calcitriol

Whole genome bisulfite sequencing reads, alignment statistics, bisulfite conversion rates and coverage rates are detailed in Additional file 1. On average, 96% of genome wide CpGs were covered at an average depth of 16x. Genome wide DNA methylation did not vary to a large extent by age or calcitriol exposure. The average proportion of methylated reads was 0.83 for each of the four sample categories (adult ± vitamin D, paediatric ± vitamin D; see Figure 2A). Multidimensional scaling analysis of methylation values by sample found little difference in DNA methylation secondary to calcitriol. Differences due to calcitriol exposure were generally much smaller than those due to individual differences or age (Figure 2B).

**Figure 2.**
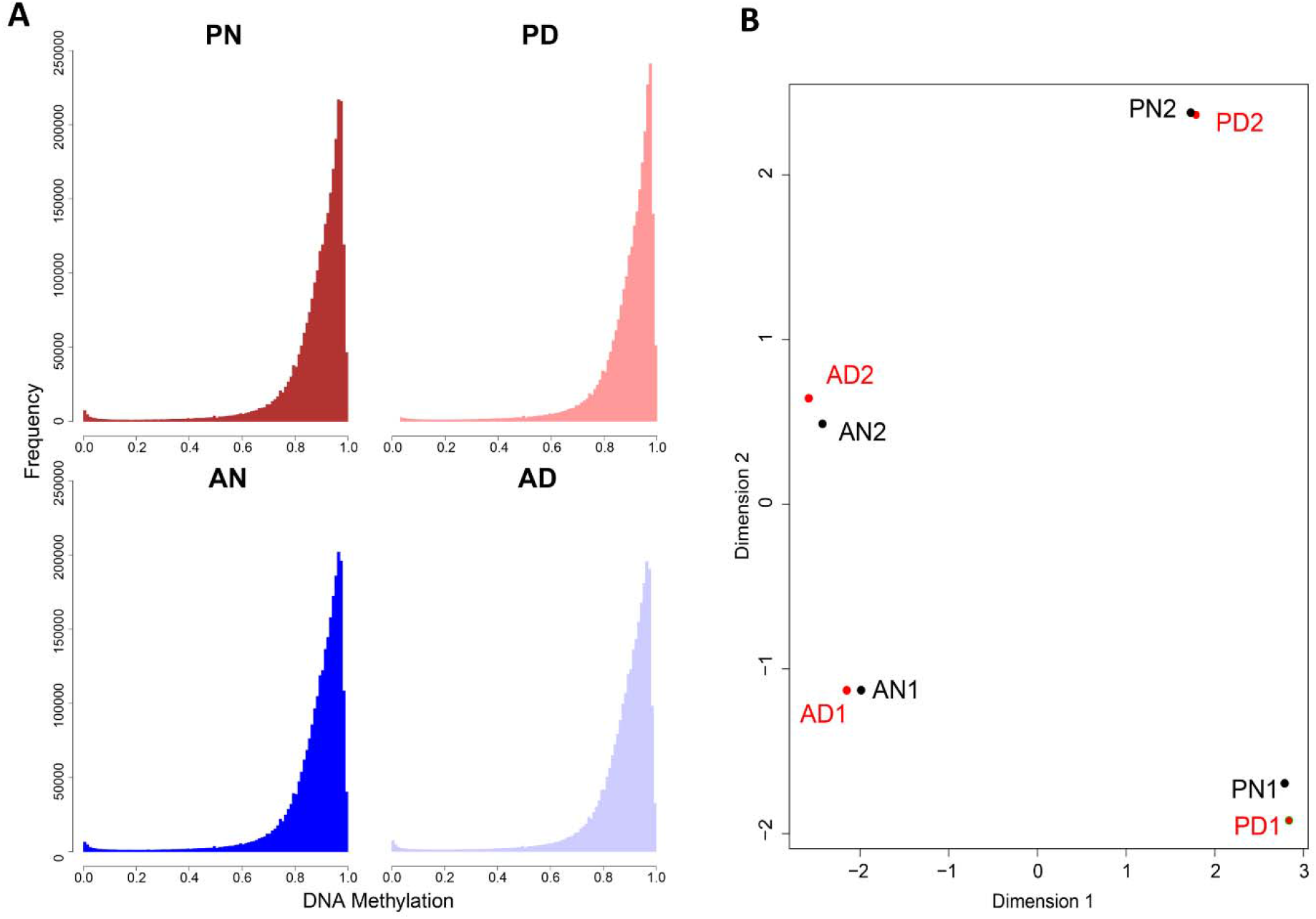
Global DNA methylation by age and calcitriol status. **A)** Frequency histograms of 1kb tile, genome wide DNA methylation, showing similar distribution of DNA methylation between conditions. **B)** Multidimensional scaling analysis of CpG wise methylation values demonstrating only minor differences in DNA methylation with the addition of calcitriol. A – adult, P – paediatric, N – no calcitriol, D – with calcitriol, 1 or 2 refer to adult or paediatric subject 1 or 2.

At an individual CpG level there were few that differed in methylation state following calcitriol exposure. In cells of adult origin, there were 382 differentially methylated CpGs (FDR<0.05) corresponding to 29 autosomal and 11 mitochondrial genes/gene promoters. Amongst cells of paediatric origin, there were 37 differentially methylated CpGs corresponding to 2 genes. None of the differentially methylated CpGs overlapped between adult and paediatric samples or with VDR peaks. Of all adult differentially methylated CpGs, a subset mapped to the promoter region of one MS risk gene, PAPD7. Details of differentially methylated CpGs can be found in Additional file 2 and 3.

### DNA Methylation Varies Markedly by Donor Age at Myeloid VDR Peaks

VDR binding sites are another mechanism by which calcitriol may exert effects on gene expression. There were marked differences in the distribution of DNA methylation between samples of adult and paediatric origin at myeloid VDR peaks, which were not apparent at other transcription factor binding sites or regulatory regions (Figure 3). There was overall lower DNA methylation at myeloid VDR peaks in cells of paediatric origin (p<2.2×10^−16^, Wilcoxon Rank Sum Test). RADmeth^20^ was used to call differentially methylated CpGs between samples of adult and paediatric origin regardless of exposure to calcitriol. There were 26134 differentially methylated CpGs corresponding to 7244 VDR peaks (52% of all myeloid VDR peaks) and 2973 genes (Additional file 4). In comparison, analysis of CD14+ transcription factor binding sites (TFBS) yielded 7125 differentially methylated CpGs corresponding to 1896 TFBS (22% of annotated CD14+ TFBS). In comparison to TFBS, differential methylation was proportionally greater at VDR peaks than TFBS (χ^2^=3983, p<1×10^−5^). A statistical overrepresentation test^21^ (Panther GO-slim annotation version 14.1, released March 12, 2019) found many immune and intracellular signalling ontologies to be enriched amongst genes corresponding to differentially methylated myeloid VDR peaks (Figure 4B).

**Figure 3.**
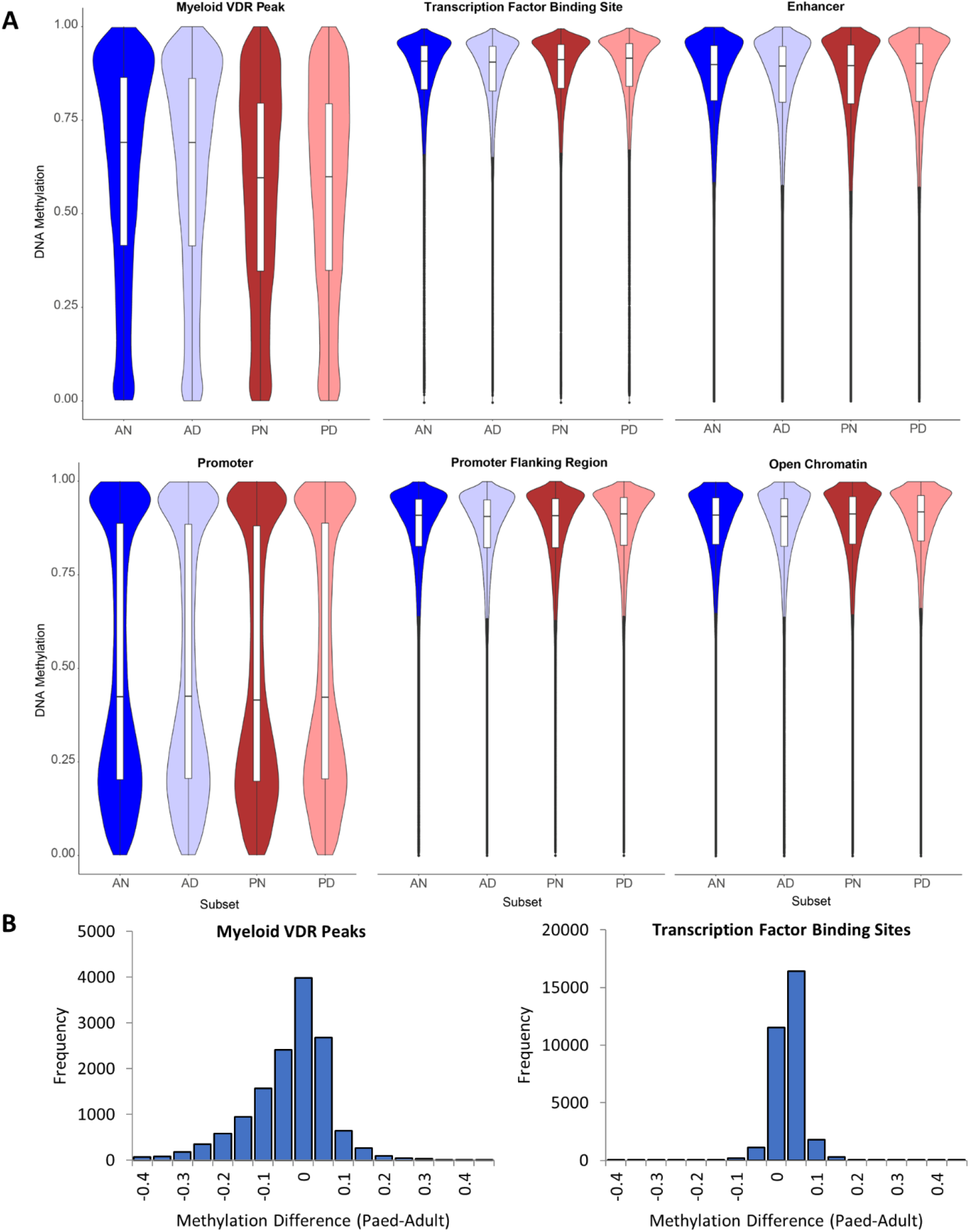
Distribution of DNA methylation by genomic feature. **A)** Violin plots of DNA methylation across various genomic features by cell origin and culture condition. Myeloid VDR peaks demonstrated DNA methylation that was skewed towards lower methylation levels in cells of paediatric origin in comparison to those of adult origin. The effects of calcitriol on DNA methylation distribution was not evident. **B)** Methylation difference between cells of paediatric and adult origin at myeloid VDR peaks and transcription factor binding sites, showing skewing towards paediatric hypomethylation at myeloid VDR peaks, but not at transcription factor binding sites. A – adult, P – paediatric, N – no calcitriol, D – with calcitriol, 1 or 2 refer to adult or paediatric subject 1 or 2.

**Figure 4.**
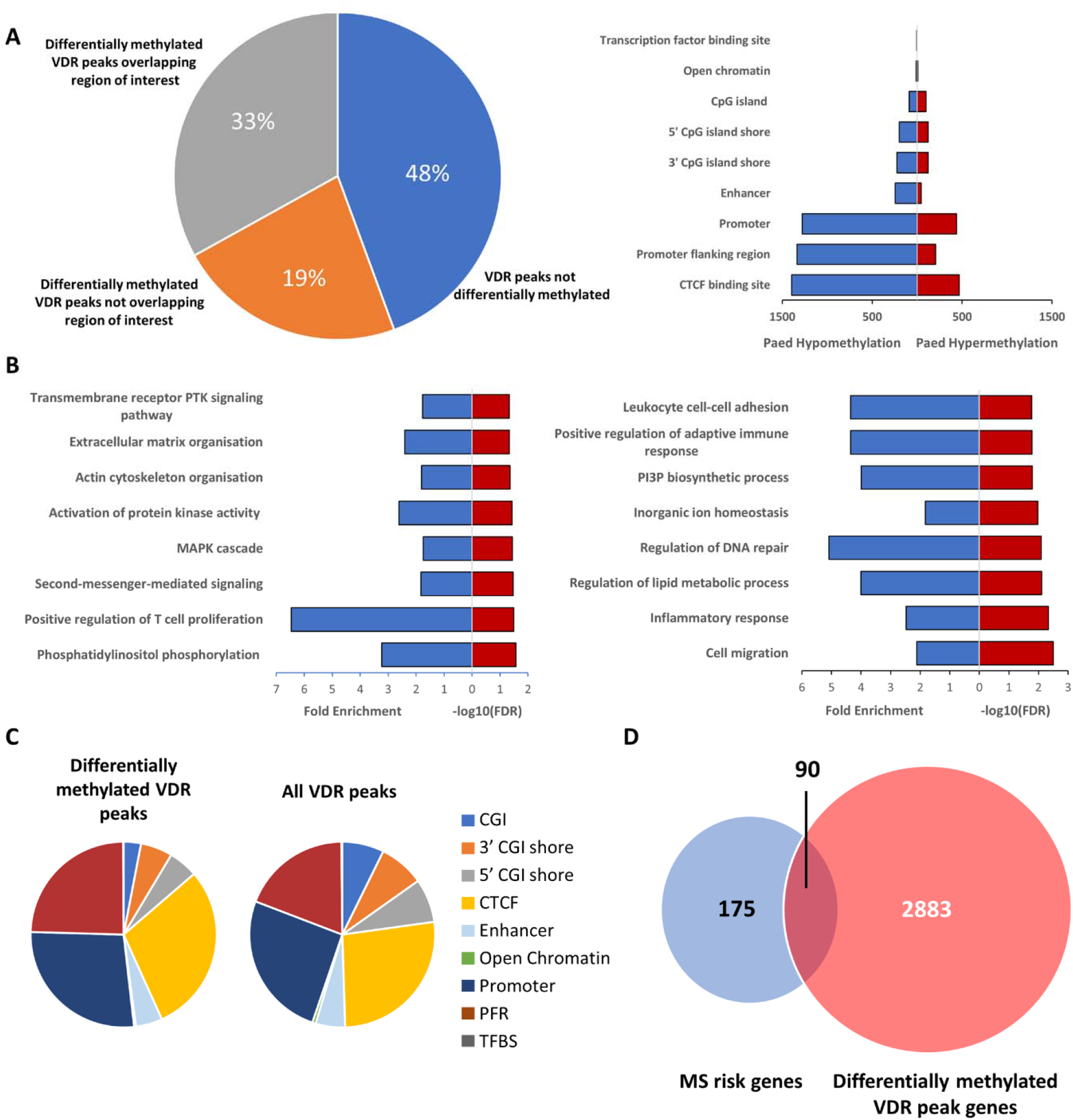
Characteristics of differentially methylated VDR peaks. **A)** Breakdown of VDR peaks based on differential methylation status and overlap with regions of interest (regulatory regions, CpG islands and island shores; left) and differentially methylated VDR peaks overlapping with regions of interest (right). The majority of regulatory regions demonstrated hypomethylation in cells of paediatric origin. **B)** Overrepresented GO terms (FDR <0.05) associated with differentially methylated VDR peaks. **C)** Breakdown of differentially methylated myeloid VDR peaks and corresponding annotation overlaps compared with all annotated VDR peaks. **D)** Overlap of currently known non-HLA MS risk genes and their overlap with differentially methylated myeloid VDR peaks. PTK *–* protein tyrosine kinase, PI3P – phosphatidylinositol-3-phosphate, CGI – CpG island, PFR – promoter flanking region, TFBS – transcription factor binding site.

The genomic annotations overlapping differentially methylated myeloid VDR peaks were then determined. Of interest, were CD14+ Ensembl regulatory build annotations and hg19 CpG island/shore annotations. Five prime and 3’ CpG island shores were designated as 2000bp upstream and downstream of the corresponding hg19 CpG island respectively. Fifty-two percent of myeloid VDR peaks contained differentially methylated CpGs, with most of these being hypomethylated in cells of paediatric origin relative to cells of adult origin. The peaks overlapped predominantly with promoter regions, promoter flanking regions and CTCF binding sites, although not in the expected proportion in comparison to all myeloid VDR peaks (χ^2^=300.8, p<2.2×10^−16^, df=8), suggestive of enrichment for specific genomic annotations (Figure 4C). The overlap of 352 CpGs previously identified as markers of biological age^22^ with differentially methylated VDR peaks was ascertained to determine whether differential methylation could be attributed to the cumulative effects of epigenetic maintenance. None of the differentially methylated VDR peaks contained a “clock” CpG. Ninety of the differentially methylated VDR peaks overlapped with non-HLA multiple sclerosis risk genes (Figure 4D), underlining the potential importance of DNA methylation differences in this latitude dependent autoimmune disease.

### Transcriptomic Effects of Calcitriol Vary by Age

Transcriptomic analysis by RNA-seq identified 183 downregulated and 154 upregulated genes amongst adult cells treated with calcitriol compared to no calcitriol, using a fold-change threshold of two. Amongst cells of paediatric origin, there were 167 downregulated and 158 upregulated genes due to calcitriol (Figure 5A). Overall, only 75 differentially expressed genes overlapped between cells of adult and paediatric origin (Figure 5B), 57 in the same direction with calcitriol exposure and 18 in opposite directions (Additional file 6). A statistical overrepresentation test did not yield any statistically significant GO terms associated with any of the differentially expressed gene sets. None of the differentially expressed genes secondary to calcitriol were differentially methylated.

**Figure 5.**
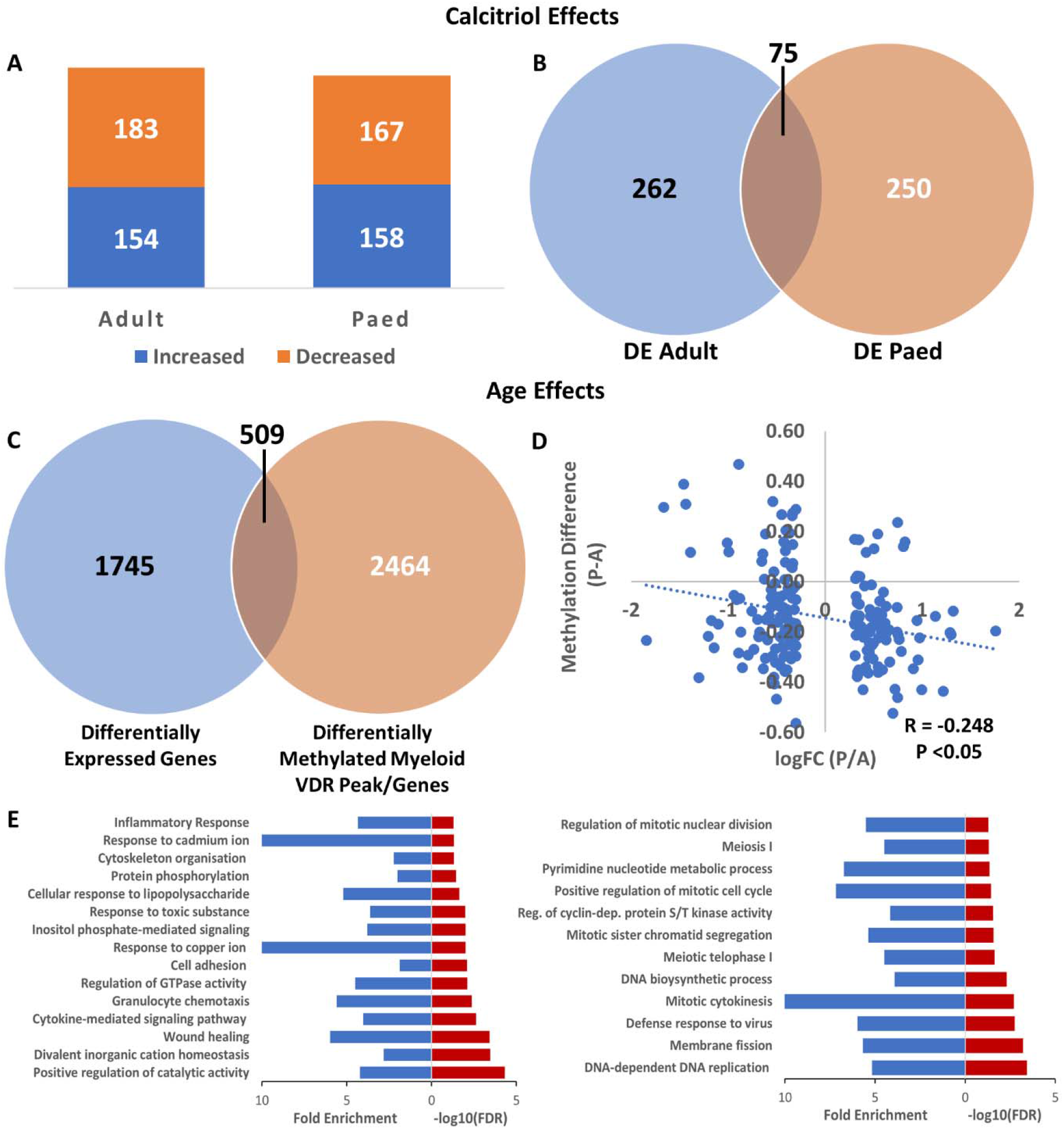
Differentially expressed genes, associated gene ontologies and DNA methylation overlap. **A)** Number of genes demonstrating changes in expression (>2 fold) due to calcitriol in cells of adult and paediatric origin. **B)** A minority of common genes are differentially expressed in response to calcitriol amongst cells of adult and paediatric origin. **C)** Overlap between differentially expressed genes and differentially methylated myeloid VDR peaks/genes when comparing cells of adult and paediatric origin. **D)** Scatter plot of overlapping sites from C) corresponding to annotated promoter regions. There was a significant negative correlation between methylation difference (paediatric – adult) and log fold-change (paediatric/adult). **E)** Overrepresented GO biological process terms (FDR<0.05) of differentially expressed sites from C) with two-fold decreased expression in cells of paediatric origin (left) or two-fold increase in cells of paediatric origin (right). DE – differentially expressed

### Age-Dependent Transcriptomic Effects Are Greater Than Calcitriol Dependent Effects

A greater number of differentially expressed genes were observed between adult and paediatric cells, independent of calcitriol exposure. We found 1002 genes were downregulated and 1252 upregulated in cells of paediatric compared to adult origin. Of these genes, 509 overlapped with differentially methylated myeloid VDR peaks (p<9.38×10^−6^, hypergeometric test; Figure 5C & 6; co-location with genomic annotations is noted in Additional file 7). There was a negative correlation between expression fold-change and methylation difference at CD14+ annotated promoters, consistent with the known relationship between DNA methylation at promoter regions and gene expression (Figure 5D). Of the differentially expressed genes, those underexpressed in cells of paediatric origin were enriched for biological processes relating to inflammation, intracellular signalling and metal ion homeostasis. Those overexpressed in cells of paediatric origin were associated with cellular replication and cell cycle processes (Figure 5E).

**Figure 6.**
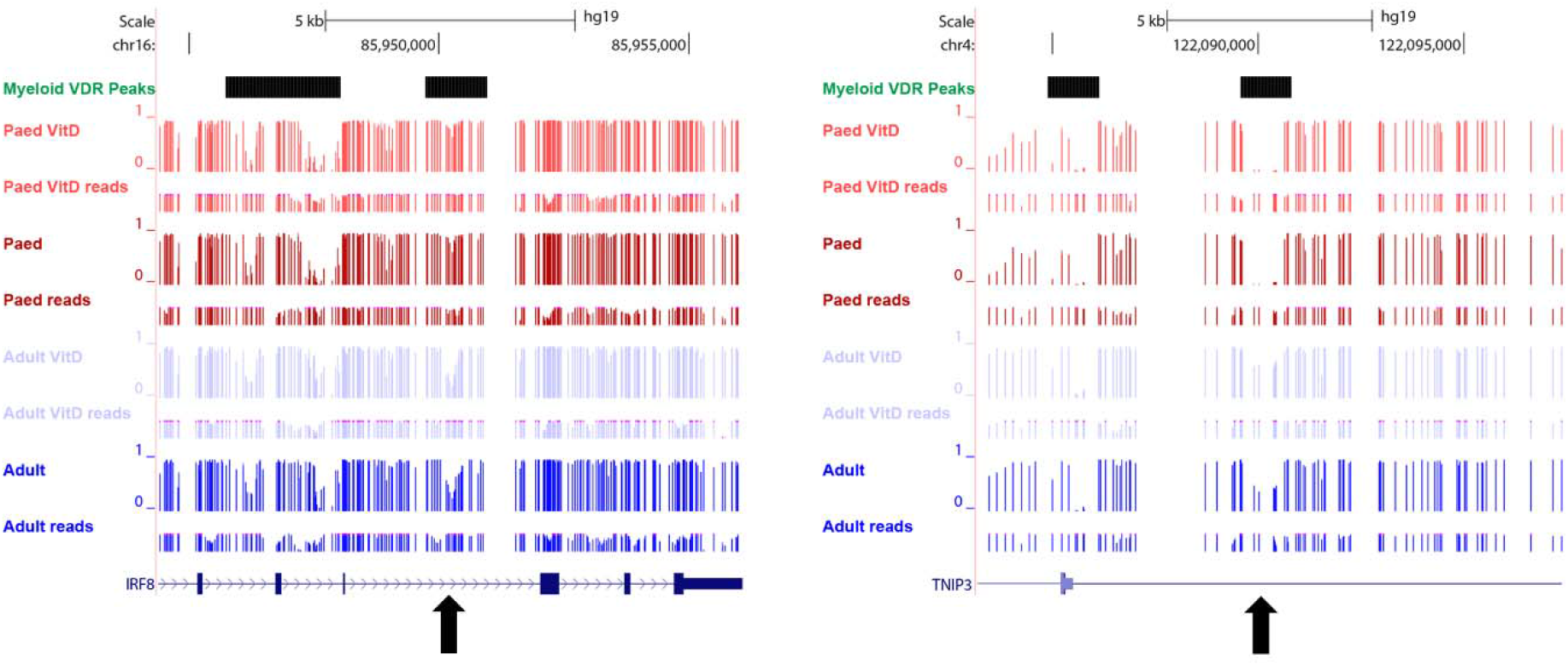
An example of differential methylation at myeloid VDR peaks overlapping with MS risk genes. In cells of paediatric origin, DNA methylation was increased at IRF8 (left) and decreased at TNIP3 (right) relative to cells of adult origin. Both genes were also differentially expressed between cells of adult and paediatric origin (see Additional file 7). Black arrows denote differentially methylated regions. Red tracks – paediatric, blue tracks – adult.

Of the 509 overlapping differentially expressed/methylated genes, 28 overlapped with 265 non-HLA MS risk genes (p=1.82×10^−6^, hypergeometric test). Two hundred and seventy-six of these overlapping genes underexpressed in paediatric cells were enriched for the GO terms “cell surface receptor signalling”, “cell migration”, “intracellular signal transduction” and “protein phosphorylation”. The remaining 233 overexpressed genes were not enriched for any biological process terms (see Additional file 7).

## Discussion

This study examined potential interactions between calcitriol and DNA methylation in myeloid cells, to identify mechanisms underlying critical periods in the development of latitude dependent autoimmune diseases such as MS. Calcitriol addition resulted in marked morphologic and phenotypic effects, however, DNA methylation changes were relatively minor in comparison. Vitamin independent DNA methylation changes differed between cells of paediatric and adult origin, especially at myeloid VDR peaks. MS risk genes were prominent among differentially methylated VDR peaks. Gene expression changes due to calcitriol, were marked, and differed strikingly between adult and paediatric cells, but only ~22% of differentially expressed genes were common to both conditions. The changes in VDR peak methylation due to age and calcitriol may be sufficient to drive the profoundly different transcriptomes between paediatric and adult derived mononuclear phagocytic cells, with evidence of MS risk gene involvement.

The calcitriol bound VDR-RXR complex binds to VDREs and participates in transcriptional regulation. Therefore, DNA methylation changes at these regions are likely to have important functional consequences. We found 52% of myeloid VDR peaks were differentially methylated between cells of adult and paediatric origin. Differential methylation was enriched above background CD14+ TFBS, providing support for VDR specificity of age-dependent changes. Most of the differentially methylated peaks also displayed decreased methylation in cells of paediatric origin. At 17% of these sites, there was concomitant differential gene expression, suggesting an immediate functional effect of these methylation differences within a subset of myeloid VDR peaks. Biological processes associated with underexpressed genes in paediatric cells were predominantly associated with inflammation, intracellular signalling and response to divalent cations, whereas those associated with overexpressed genes were predominantly associated with cellular proliferation.

Genes in *cis* with most differentially methylated VDR peaks were associated with biological processes important in inflammation and cellular differentiation, but not with changes in gene expression. VDR binding sites proximal to genes encoding PI3K subunits (including PIK3R1/3/6, PIK3CG, PIK3C2B) were differentially methylated between cells of adult and paediatric origin. The PI3K molecular pathway plays a role in myeloid cell differentiation^23,24^, monocyte antimycobacterial activity^25^ and promotion of macrophage differentiation^26^. Genes relating to the MAPK cascade were also enriched amongst differentially methylated VDR peak genes. The MAPK cascade is involved in significant crosstalk with the PI3K/AKT pathway^27^ and has pleiotropic effects in monocytes/macrophages depending on the triggering stimulus and cell type. These effects include differentiation and activation^28^. Together, the differential methylation of genes relating to both the PI3K and MAPK pathways suggest that the differing potential for vitamin D related myeloid cell differentiation between adult and paediatric cells may be encoded by DNA methylation.

Genes belonging to ontologies relating to the regulation of the adaptive immune response and regulation of T cell proliferation were also associated with differential methylation between adult and paediatric cells. This suggests that myeloid cells are differentially primed to influence the adaptive immune system in childhood compared with adulthood.

Consistent with previous work on vitamin D supplementation in mononuclear cells^19^, DNA methylation in our myeloid cells appeared to be relatively insensitive to the effects of calcitriol. This occurred despite differentiation from haematopoietic progenitor cells from paediatric donors, which would presumably demonstrate greater DNA methylation plasticity in response to environmental stimuli. In contrast, the previously documented effect of vitamin D on DNA methylation in mouse CD4+ T cells was much more prominent^14^. Many effects of vitamin D on human monocytes/macrophages may be mediated by epigenetic marks other than DNA methylation^18^.

How age-dependent methylomic differences at myeloid VDR binding sites confer long term risk for latitude dependent diseases is unclear. Monocytes typically persist in the circulation for up to ~1 week^29^, and would be an unlikely substrate for DNA methylation dependent risk unless they migrate to peripheral sites. These tissue resident macrophages are known to persist for much longer periods (months to years)^30^, perhaps transmitting early life influenced phenotypic changes that either predispose or limit propensity to autoimmunity in later life (Figure 7). Another possibility is that VDR agonism (or lack thereof) during early life leads to persistent changes in VDR binding site methylation and later susceptibility to VDR effector activities.

**Figure 7.**
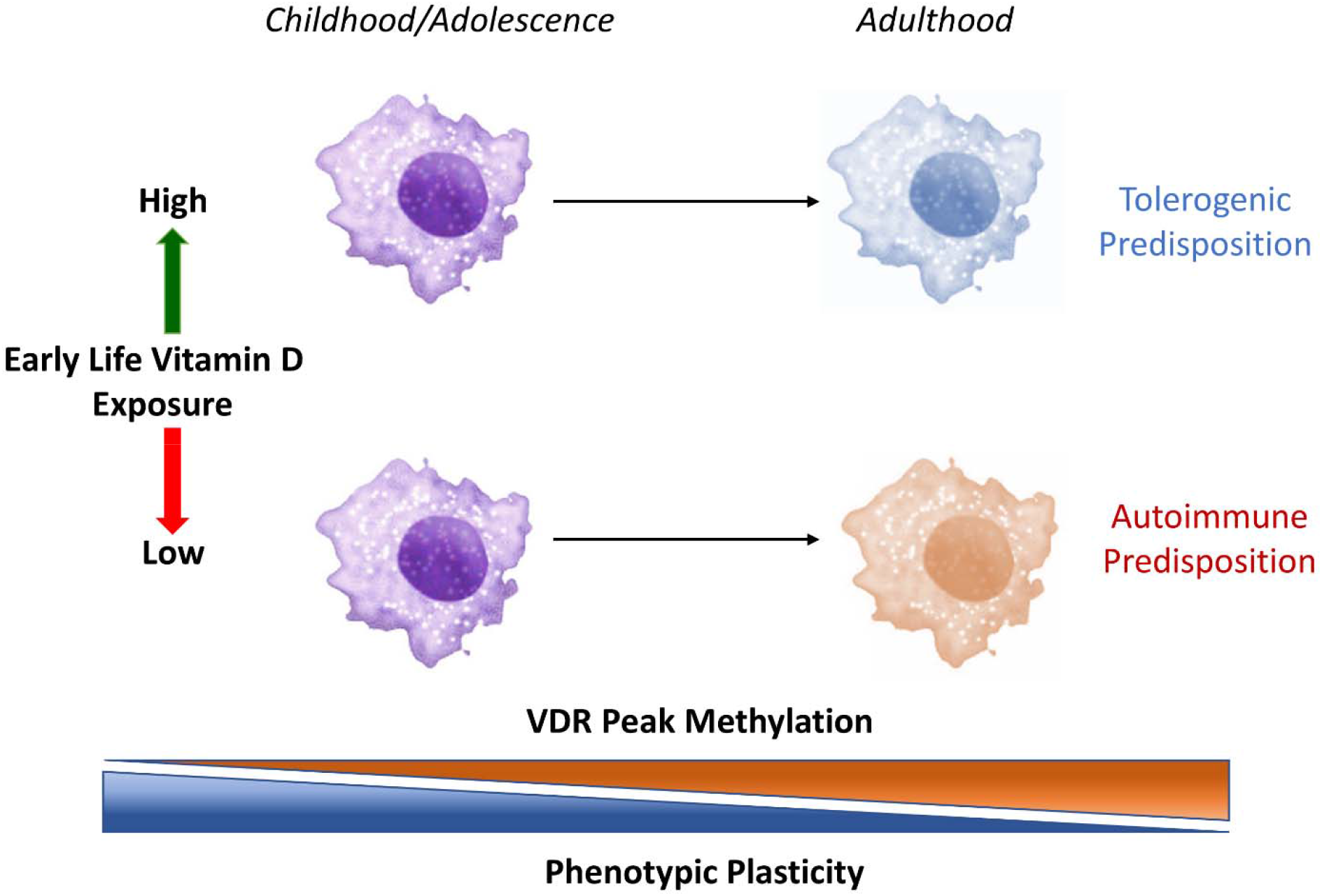
A potential mechanism for the development of autoimmune disease risk dependent on early life vitamin D exposure. Decreased VDR binding site methylation in early life increases phenotypic plasticity and susceptibility to vitamin D exposure. Because tissue macrophages persist for months to years, phenotypic settings resulting from vitamin D exposure in early life may lead to a tolerogenic or autoimmune propensity in later life. Macrophage images adapted from^31^.

One potential reason for finding minimal DNA methylation changes with vitamin D could be related to the duration of the cell cultures. Increasing passage number is associated with increases in DNA methylation^22^, potentially distorting or obscuring the effects of vitamin D or age on DNA methylation. Due to the lack of CD14+ cells at earlier stages of culture, we were also unable to determine temporal effects on DNA methylation with vitamin D exposure. It has been previously shown that chromatin accessibility due to calcitriol exposure peaks at 24 hours and virtually returns to baseline levels after 48 hours^18^. It could be argued however, that DNA methylation changes are less likely to occur within these time frames in comparison to histone modifications and non-coding RNAs. Finally, whether the phenomenon of age-related differential methylation at VDR binding sites occurs *in vivo* requires further investigation.

Future studies will need to ascertain the robustness of our present findings across a greater number of biological replicates. This study also raises questions regarding age-dependent VDR methylation in other cell lineages as well as haematopoietic progenitor cells, and whether VDR methylation settings might be transmitted to progeny cells. The model of altered tissue macrophage phenotype might also be amenable to study by comparison of their phenotype/function in MS with normal individuals, for example in co-culture.

## Conclusions

Whilst vitamin D has minor effects on the myeloid methylome, age-dependent differences in VDR peak DNA methylation suggest vitamin D exposure at critical periods in immune system development may contribute to well characterised latitude related differences in autoimmune disease risk.

## Declarations

### Ethics approval and consent to participate

This study received ethics approval from the Western Sydney Local Health District Human Research Ethics Committee (HREC2002/9/3.6(1425) & (5366) AU RED LNR/17/WMEAD/447).

### Consent for publication

Not applicable

### Availability of data and materials

The datasets used and/or analysed during the current study are available from the corresponding author on reasonable request.

### Authors’ Contributions

LO, GP and DB devised the experiments. NF and SS assisted in planning and analysis of cell culture and flow cytometric experiments. LO conducted the experiments, analyses and prepared the manuscript. GP performed RNA-seq and assisted in data analysis. All authors read and approved the final manuscript.

## Acknowledgements

The authors would like to acknowledge Australian Red Cross Blood Services, the Sydney Cord Blood Bank and donors for providing samples for this study. The authors also acknowledge Prof David Brown for his invaluable comments on the manuscript. Flow cytometry was performed at the Flow Cytometry Core Facility supported by the Westmead Research Hub, Cancer Institute NSW and NHMRC. Bioinformatic analysis was supported by Sydney Informatics Hub, funded by the University of Sydney.

## Additional Files

Additional file 1 – Cell harvest, WGBS alignment, bisulfite conversion and coverage statistics; Additional file 1.xlsx

Additional file 2 – Genome wide paediatric differentially methylated CpGs; Additional file 2.xlsx

Additional file 3 – Genome wide adult differentially methylated CpGs; Additional file 3.xlsx

Additional file 4 – Differentially methylated VDR myeloid peaks; Additional file 4.xlsx

Additional file 5 – MS risk genes overlapping differentially methylated VDR peaks; Additional file 5.xlsx

Additional file 6 – Differentially expressed genes, vitamin D vs no vitamin D; Additional file 6.xlsx

Additional file 7 – Differentially expressed genes, adult vs paediatric; Additional file 7.xlsx

## References

1. Wacker M, Holick MF. Sunlight and Vitamin D: A global perspective for health. Dermato-endocrinology 2013; 5(1): 51–108.

2. Osborne NJ, Ukoumunne OC, Wake M, Allen KJ. Prevalence of eczema and food allergy is associated with latitude in Australia. Journal of allergy and clinical immunology 2012; 129(3): 865–7.

3. Heim C, Binder EB. Current research trends in early life stress and depression: Review of human studies on sensitive periods, gene–environment interactions, and epigenetics. Experimental neurology 2012; 233(1): 102–11.

4. Cedar H, Bergman Y. Linking DNA methylation and histone modification: patterns and paradigms. Nature Reviews Genetics 2009; 10(5): 295–304.

5. Ahlgren C, Lycke J, Odén A, Andersen O. High risk of MS in Iranian immigrants in Gothenburg, Sweden. Multiple sclerosis journal 2010; 16(9): 1079–82.

6. Gale CR, Martyn CN. Migrant studies in multiple sclerosis. Progress in neurobiology 1995; 47(4-5): 425–48.

7. Ahlgren C, Odén A, Lycke J. A nationwide survey of the prevalence of multiple sclerosis in immigrant populations of Sweden. Multiple Sclerosis Journal 2012; 18(8): 1099–107.

8. Acevedo N, Reinius LE, Vitezic M, et al. Age-associated DNA methylation changes in immune genes, histone modifiers and chromatin remodeling factors within 5 years after birth in human blood leukocytes. Clinical epigenetics 2015; 7(1): 34.

9. Alisch RS, Barwick BG, Chopra P, et al. Age-associated DNA methylation in pediatric populations. Genome research 2012: gr. 125187.111.

10. Tobi EW, Lumey L, Talens RP, et al. DNA methylation differences after exposure to prenatal famine are common and timing-and sex-specific. Human molecular genetics 2009; 18(21): 4046–53.

11. Tobi EW, Slieker RC, Stein AD, et al. Early gestation as the critical time-window for changes in the prenatal environment to affect the adult human blood methylome. International journal of epidemiology 2015; 44(4): 1211–23.

12. Anderson CM, Gillespie SL, Thiele DK, Ralph JL, Ohm JE. Effects of Maternal Vitamin D Supplementation on the Maternal and Infant Epigenome. Breastfeeding medicine : the official journal of the Academy of Breastfeeding Medicine 2018; 13(5): 371–80.

13. Zhu H, Bhagatwala J, Huang Y, et al. Race/ethnicity-specific association of vitamin D and global DNA methylation: cross-sectional and interventional findings. PloS one 2016; 11(4): e0152849.

14. Zeitelhofer M, Adzemovic MZ, Gomez-Cabrero D, et al. Functional genomics analysis of vitamin D effects on CD4+ T cells in vivo in experimental autoimmune encephalomyelitis. Proceedings of the National Academy of Sciences of the United States of America 2017; 114(9): E1678–e87.

15. Moore JR, Hubler SL, Nelson CD, Nashold FE, Spanier JA, Hayes CE. 1,25-Dihydroxyvitamin D3 increases the methionine cycle, CD4(+) T cell DNA methylation and Helios(+)Foxp3(+) T regulatory cells to reverse autoimmune neurodegenerative disease. J Neuroimmunol 2018; 324: 100–14.

16. Parnell GP, Booth DR. The multiple sclerosis (MS) genetic risk factors indicate both acquired and innate immune cell subsets contribute to MS pathogenesis and identify novel therapeutic opportunities. Frontiers in immunology 2017; 8: 425.

17. Shahijanian F, Parnell GP, McKay FC, et al. The CYP27B1 variant associated with an increased risk of autoimmune disease is underexpressed in tolerizing dendritic cells. Human molecular genetics 2014; 23(6): 1425–34.

18. Seuter S, Neme A, Carlberg C. Epigenome-wide effects of vitamin D and their impact on the transcriptome of human monocytes involve CTCF. Nucleic acids research 2016; 44(9): 4090–104.

19. Chavez Valencia RA, Martino DJ, Saffery R, Ellis JA. In vitro exposure of human blood mononuclear cells to active vitamin D does not induce substantial change to DNA methylation on a genome-scale. The Journal of steroid biochemistry and molecular biology 2014; 141: 144–9.

20. Dolzhenko E, Smith AD. Using beta-binomial regression for high-precision differential methylation analysis in multifactor whole-genome bisulfite sequencing experiments. BMC bioinformatics 2014; 15(1): 215.

21. Mi H, Huang X, Muruganujan A, et al. PANTHER version 11: expanded annotation data from Gene Ontology and Reactome pathways, and data analysis tool enhancements. Nucleic acids research 2016; 45(D1): D183–D9.

22. Horvath S. DNA methylation age of human tissues and cell types. Genome Biology 2013; 14(10): 3156.

23. Hmama Z, Nandan D, Sly L, Knutson KL, Herrera-Velit P, Reiner NE. 1α, 25-dihydroxyvitamin D3–induced myeloid cell differentiation is regulated by a vitamin D receptor–phosphatidylinositol 3-kinase signaling complex. The Journal of experimental medicine 1999; 190(11): 1583–94.

24. Neri LM, Marchisio M, Colamussi ML, Bertagnolo V. Monocytic differentiation of HL-60 cells is characterized by the nuclear translocation of phosphatidylinositol 3-kinase and of definite phosphatidylinositol-specific phospholipase C isoforms. Biochemical and biophysical research communications 1999; 259(2): 314–20.

25. Sly LM, Lopez M, Nauseef WM, Reiner NE. 1α, 25-Dihydroxyvitamin D3-induced monocyte antimycobacterial activity is regulated by phosphatidylinositol 3-kinase and mediated by the NADPH-dependent phagocyte oxidase. Journal of Biological Chemistry 2001; 276(38): 35482–93.

26. Liu Q, Ning W, Dantzer R, Freund GG, Kelley KW. Activation of protein kinase C-ζ and phosphatidylinositol 3′-kinase and promotion of macrophage differentiation by insulin-like growth factor-I. The Journal of Immunology 1998; 160(3): 1393–401.

27. Aksamitiene E, Kiyatkin A, Kholodenko BN. Cross-talk between mitogenic Ras/MAPK and survival PI3K/Akt pathways: a fine balance. Portland Press Ltd.; 2012.

28. Rao KMK. MAP kinase activation in macrophages. Journal of leukocyte biology 2001; 69(1): 3–10.

29. Patel AA, Zhang Y, Fullerton JN, et al. The fate and lifespan of human monocyte subsets in steady state and systemic inflammation. Journal of Experimental Medicine 2017; 214(7): 1913–23.

30. Parihar A, Eubank TD, Doseff AI. Monocytes and macrophages regulate immunity through dynamic networks of survival and cell death. J Innate Immun 2010; 2(3): 204–15.

31. Taylor C. 3 CELLS fvcc104. OpenStax CNX 3 Nov 2019. http://cnx.org/contents/21a91101-9df0-4826-bb9b-94883dcffcc4@1.1.

